# Enhancing NADPH to Restore Redox Homeostasis and Lysosomal Function in G6PD-Deficient Microglia

**DOI:** 10.1101/2024.09.12.607918

**Authors:** Abir Mondal, Soumyadeep Mukherjee, Prince Upadhyay, Isha Saxena, Soumya Pati, Shailja Singh

## Abstract

Microglia, the residential immune cells in the central nervous system (CNS) exhibited in multiple states from resting to activated, play a significant role in neurogenesis, myelination, synaptic transmission, immune surveillance, and neuroinflammation. The aggravated inflammatory response by microglia triggers the generation of superoxide, which often causes the degeneration of neurons, leading to the development of Parkinson’s and Alzheimer’s. The oxidative stress is key to many neurological disorders, often regulated by many genes. The terminal neutralization of oxidative stress is mediated by NADPH and glutathione. The cytosolic NADPH level is majorly contributed by a key enzyme called glucose-6-phosphate dehydrogenase (G6PD). The deficiency of G6PD is associated with hemolytic anemia, diabetes, cardiovascular, autoimmune, and neurological disorders. Our recent study indicated that G6PD deficiency decreases cytosolic NADPH levels and alters redox homeostasis and lysosomal function in microglia. Therefore, replenishment of NADPH is crucial for targeting G6PD deficiency-mediated microglial dysfunctions. This research promotes alternate metabolic pathways by targeting the expression and activity of enzymes such as isocitrate dehydrogenase 1 (IDH1) and malic enzyme 1 (ME1), which are responsible for cytoplasmic NADPH production. Metabolites like citric and malic acid supplementation promote NADPH production and reduce microglial oxidative stress. Additionally, using another group of small-molecule metabolites, such as dieckol and resveratrol, enhances the expression of IDH1 and ME1 enzymes to resolve potential tissue heterogeneity. Finally, combining these metabolites supplementation increased NADPH production and restored redox homeostasis and lysosomal function in G6PD deficient microglia, indicating their further use as potential therapeutics against G6PD deficiency disorders.

**Graphical abstract:** 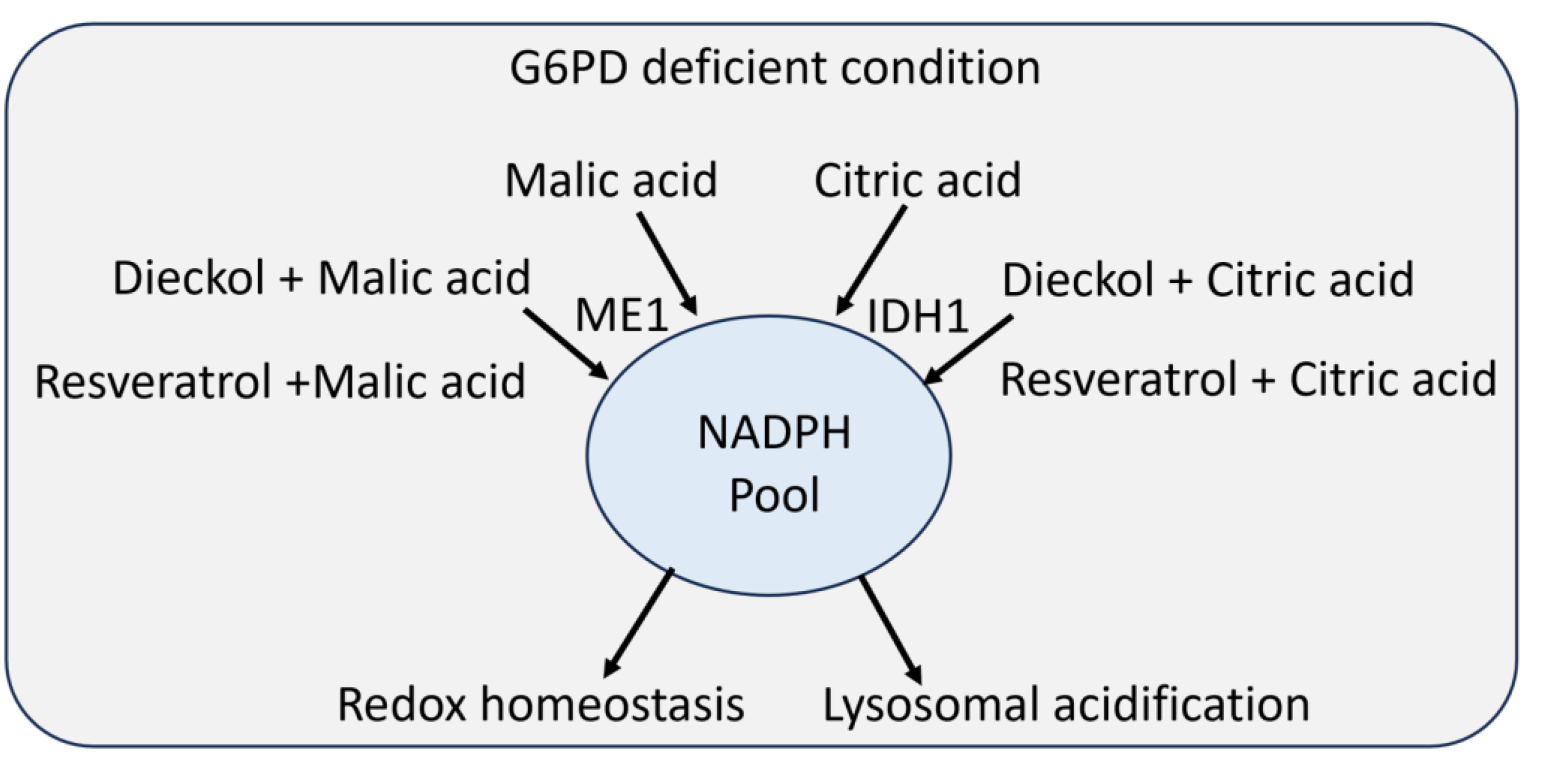

## 1. Introduction

Microglia plays a crucial role in pathogenic invasion and injury and maintains the physiological function of the CNS, particularly under oxidative stress. [1]. Oxidative stress occurs due to the imbalance between reactive oxygen species (ROS) production and the cellular antioxidant defenses. Microglia are deeply involved in both the production and mitigation of oxidative stress, highlighting their dual role in CNS health and disease [2]. The chronic activation of microglia can exacerbate neuronal injury and death. On the contrary, microglia are essential for neurogenesis, myelination, synapse modulation, and vasculogenesis [3]. It also plays a critical role in removing dead cells and pathogenic β-amyloid and tau plaques through phagocytosis [4]. Thus, the function of microglia is key to CNS homeostasis, and its alteration results in the pathogenesis and progression of neurodegenerative diseases, highlighting their importance as potential therapeutic targets for promoting CNS health. Microglia become activated in response to pathogens, injury, or other stress signals and produce ROS as part of the innate immune response. NADPH oxidase and mitochondrial pathways are primarily responsible for ROS production and are essential for defense against pathogens and signaling processes. However, the accumulation of ROS can lead to oxidative damage to lipids, proteins, and DNA and further promote neurodegenerative processes [5]. Microglia also produce antioxidant molecules such as NADPH and glutathione to neutralize oxidative stress, thus protecting neurons and other glial cells from oxidative damage.

However, NADPH is the key molecule that allows microglia to mediate pro and anti-oxidative functions depending on the cellular microenvironment [6]. During pathogen invasion, NADPH is being utilized to generate ROS to neutralize foreign particles. On the contrary, NADPH is crucial for detoxifying hydrogen peroxide (H_2_O_2_) in a glutathione-dependent manner. The G6PD is a key enzyme for pentose phosphate pathways, contributing to the major production of cytoplasmic NADPH. The deficiency of G6PD causes dysregulation of redox homeostasis and accelerated production of reactive oxygen species (ROS), causing damage to the cell. Notably, the decrease in cellular NADPH and dysregulation of the redox microenvironment due to the deficiency of the G6PD enzyme results in hemolytic anemia [7,8]. Several reports suggested that the G6PD gene is highly susceptible to the mutation as it contains many GC-rich sequences [9]. Besides, the mutation in this gene showed diverse pathophysiology ranging from severe pathophysiology like hemolysis to asymptomatic unless there is any oxidative stress (caused by food, drug, or infection). However, several studies indicated that oxidative stress and neuroinflammation are correlated with neurodegenerative disorders [10– 13]. Few recent studies in an animal model and postmortem human brain neurodegenerative (Alzheimer’s, Parkinson’s, and Huntington) tissue showed either reduced G6PD expression or activity is the primary cause of pathophysiology [14–16]. Interestingly, ferroptosis, iron-dependent cellular death pathways, is very common in all neurodegenerative disorders [17– 20]. A recent study also showed that cellular NADPH concentration is a key determining factor for ferroptosis [21,22]. A low level of cellular NADPH promotes ferroptosis, which might correlate with G6PD deficiency. Besides, G6PD deficiency was also reported in 2 autistic male children in Saudi Arabia [23]. Recently, a few reports suggested that an altered redox microenvironment is one of the causes of neurodevelopmental disorders such as ADHD and ASD [24–27]. Considering the uprising population of neurological cases associated with oxidative stress-induced neuroinflammation in neurological disorders indicating their possible link with G6PD deficiency. Notably, the spectrum of G6PD deficiency disorder and the unavailability of potential therapeutics put millions of individuals at life risk. In this study, we targeted the NADPH-producing alternate pathways catalyzed by IDH1 and ME1 to restore the cellular NADPH pool, further allowing microglia to regulate cellular oxidative stress and lysosomal functions.

## 2. Materials and Methods

### 2.1 Culturing of Human microglia

Human microglia clone 3 (HMC3) cells from ATCC (CRL-3304) were cultured and passaged in complete EMEM media (EMEM, 10% FBS, 1 mM Sodium Pyruvate, 1 x NEAA, and 0.1% Penicillin-streptomycin). Microglia 5 to 16 passages were used for all the experiments. Additionally, previously prepared and characterized G6PDd-deficient microglia were used in this study [28].

### 2.2 Cytotoxicity assays

HMC3 Cells were seeded at a density of 4000 cells/well in 96-well plates. The following day, varying concentrations (0 to 100μg/ml) of Dieckol and (0 to 100μM) resveratrol were prepared in complete EMEM (cEMEM) and added to the respective well. DMSO control was also taken to nullify the solvent toxicity. Plates were incubated in a 37°C incubator for 24 hours. Then, the media was removed from the cells, and 0.5 mg/ml MTT (3-(4,5-dimethylthiazol-2-yl)-2,5-diphenyl-2H-tetrazolium bromide) solution in 1x phosphate buffer saline (PBS) was added to each well for 3 hours. Then, MTT was removed, and the formazan crystal was dissolved in a 200 μl DMSO (SRL #24075) solution. Then, the plates were incubated in a 37°C incubator for 30 minutes. The absorbance at 570nm was measured in a spectrophotometer (BioTek, Synergy H1).

### 2.3 NADPH Estimation assay

NADPH level was measured by using highly sensitive WST-8 (# HY-D0831, MCE) and 1-mPMS (# HY-D0937, MCE) assay [29], and the following conditions were used (Table 1). Spectrophotometry was performed to measure absorbance at 460nm (BioTek, Synergy H1). All the experiments and replicas were further used to calculate the standard error mean (SEM). Experimental data was represented as a bar plot generated by GraphPad.

**Table 1:**
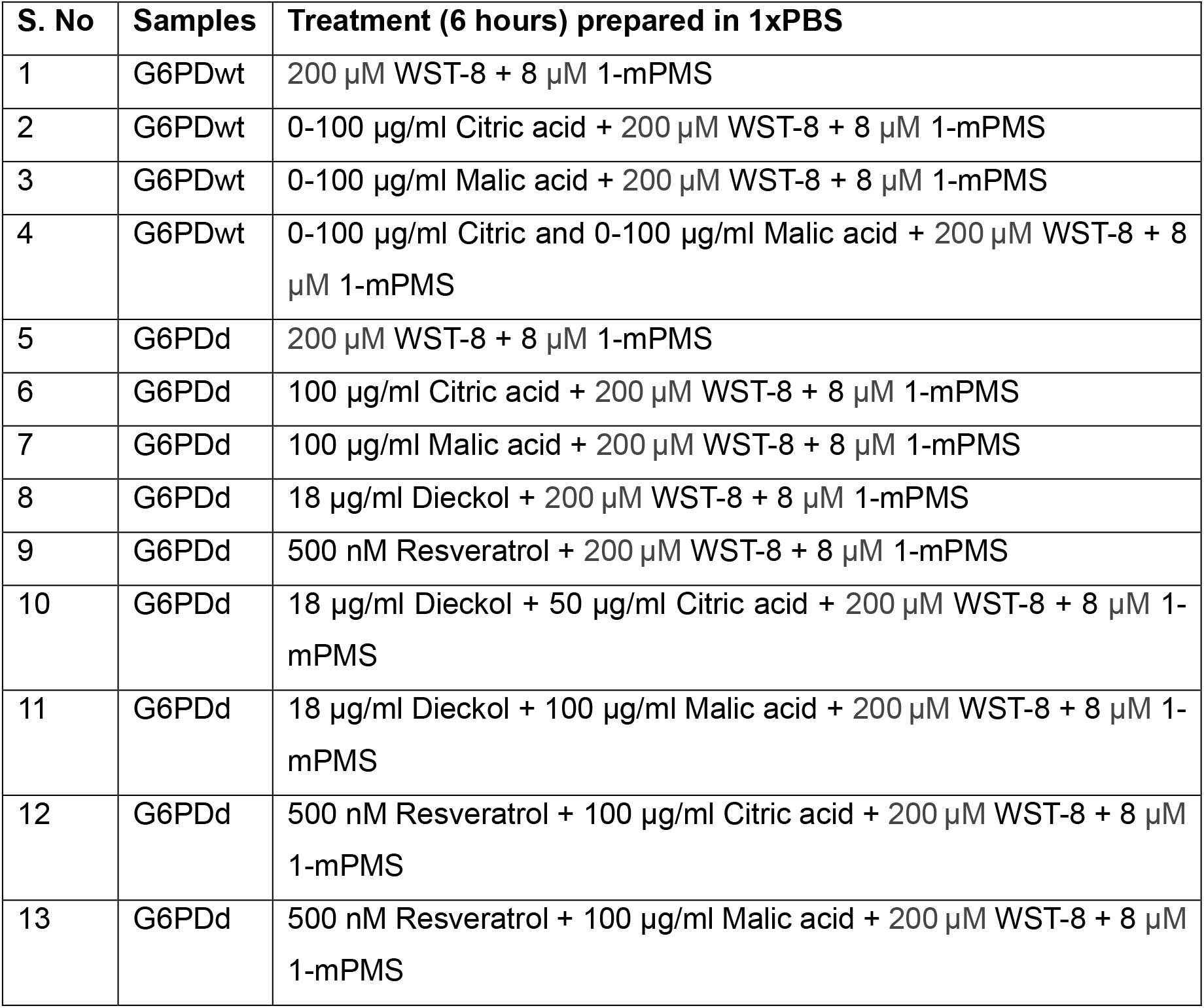
NADPH estimation by WST-8 and 1-mPMS assay.

### 2.4 Immunocytochemistry

HMC3 cells were seeded at a density of 25000 cells/coverslips. The next day, cells were treated with 0.5μM of Resveratrol and 10μg/ml Dieckol for 24 hours. Then, cells were fixed with 4% Paraformaldehyde for 10 mins and washed using 1x PBS. After that, 0.1% Triton X-100 was used to permeabilize the membrane, and blocking was performed by 5% BSA for 1 hour. Then, the NRF2 primary antibody (Affinity, Cat No. BF8017) was incubated (1:100 dilution) for 3 hours at room temperature in a humid chamber. After washing three times with 1x PBS, the anti-rabbit secondary-FITC (Bio-RAD #STAR34B) secondary antibody (1:200 dilution) was incubated for 90 mins. Then, coverslips were washed using 1x PBS and mounted on the slide using Vectashield antifade DAPI (Vector Lab, H-1200) mounting media. Images were captured using a Nikon Ti2 Confocal microscope and analyzed by Fiji ImageJ software.

### 2.5 Study the ROS by flow cytometer

1.0 × 10^5^ cells/ well were seeded in 6 well tissue culture plates. The following day, treatment was given to cells, as shown in Table 2. After the treatment, cells were trypsinized and collected by centrifugation (Eppendorf # 5810R) at 130g for 5 mins. The cell pellet was washed with 1x PBS two times to remove the residual cell culture media. Finally, the cell pellet was dissolved in 200 μl of 1x PBS. In one set, 30μM H2O2 was added to the cell and incubated for 20 minutes (positive control of our experiments). Then, 10μM carboxy-H_2_DCFDA (Invitrogen #I36007) ROS labelling dye was added to all samples incubated in a 37°C incubator for 30 mins. In one set, cells were kept without adding any dye to check the background noise. Flow cytometry was performed in CytoFLEX-S (Beckman), and 10000 events were recorded for analysis. Flow cytometry data was analyzed in CytExpert and Floreada.io online tool.

**Table 2:** Experimental conditions for detection of ROS positive cells flowcytometry.

| S No. | Sample | Pretreatment (12 hours) | Treatment (6hours) |
| --- | --- | --- | --- |
| 1 | G6PDwt |  | No dye control |
| 2 | G6PDwt |  | Control |
| 3 | G6PDwt |  | 500ng/ml LPS |
| 4 | G6PDwt | 500ng/ml LPS | 100µg/ml Citrate |
| 5 | G6PDwt | 500ng/ml LPS | 100µg/ml Malate |
| 6 | G6PDd |  | control |
| 7 | G6PDd | 500ng/ml LPS |  |
| 8 | G6PDd | 500ng/ml LPS | 100µg/ml Citrate |
| 9 | G6PDd | 500ng/ml LPS | 100µg/ml Malate |
| 10 | G6PDd |  | 100µg/ml Citrate |
| 11 | G6PDd |  | 100µg/ml Malate |
| 12 | G6PDd | 18µg/ml Dieckol |  |
| 13 | G6PDd | 500nM Resveratrol |  |
| 14 | G6PDd | 18µg/ml Dieckol | 100µg/ml Citrate |
| 15 | G6PDd | 18µg/ml Dieckol | 100µg/ml Malate |
| 16 | G6PDd | 500nM Resveratrol | 100µg/ml Citrate |
| 17 | G6PDd | 500nM Resveratrol | 100µg/ml Malate |

### 2.6 Q-PCR-based gene expression study

The RNA was isolated using a TRIZOL reagent (Thermofisher part#15596206). The RNA concentration was measured in the Nanodrop 2000 instrument (Thermo Scientific). Using a High-Capacity cDNA reverse transcription kit, 1μg of RNA was used to make complementary DNA (cDNA) (Applied Biosystems #4368814). Then, quantitative PCR (qPCR) was performed using the Powerup SYBR green master mix (Applied Biosystems # A25742). Fold change (2^-ΔΔC^t) was calculated in Microsoft Excel. The qPCR primers for respective genes are mentioned in Table 3.

**Table 3:**
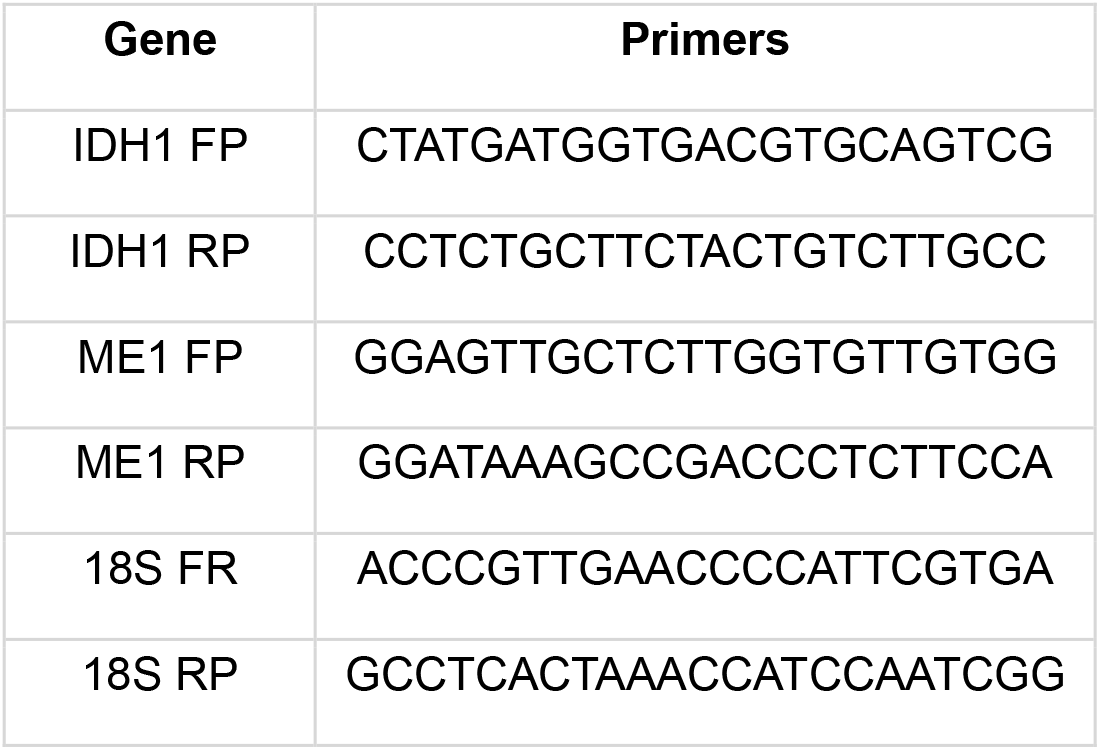
Q-PCR primer details.

### 2.7 Lysosome acidification study by confocal microscope

G6PDd microglia at a 30000 cells/ well density were seeded on a 35mm glass bottom dish. The following day, all metabolites and combination therapeutics were supplemented to cells for 6 hours in cEMEM media. Only the NADPH was incubated for 30mins which acts as a positive control for our experiments. Then, cells were labelled with 70 nM LysoTracker Deep Red (Invitrogen #L12492) and NucBlue Live Cell Stain (Hoechst 33342, R37605 for 20 minutes. Then, the probe was washed, and the fresh cEMEM was added. After that, confocal imaging was performed in a Nikon Ti2 confocal microscope. The acidified lysosomal bright puncta were counted manually on Fiji-image J software.

### 2.8 Statistical analysis

All the experiments were performed more than thrice with multiple replicates. GraphPad Prism was used to calculate the standard error mean (SEM). Statistical significances were calculated using either t-test and one-way or two-way ANOVA based on experimental parameters. P-values and statistical tests were mentioned in the respective figure legends.

## 3. Results

### 3.1 Analysis of the expression of NADPH-producing enzymes and formulation of metabolic supplementation strategies

The pentose phosphate pathway produces 2 moles of NADPH from 1 mole of Glucose-6-phosphate, which is a major contributor to cytosolic NADPH. The G6PD is the key rate-limiting enzyme of PPP, which catalyses the first step of NADPH production, followed by 6-phosphogluconate dehydrogenase (6PGD) (Figure 1A). Alternatively, isocitrate dehydrogenase (IDH1) and Malic enzyme 1 (ME1) also contribute to 1 mole of NADPH production in a tissue-specific manner (Figure 1 A). Therefore, understanding the expression of NADPH-producing genes is crucial for developing metabolic supplementation strategies. Here, we analyzed the expression of G6PD, IDH1, and ME1 in the brain using the Brainspan RNA sequencing database [30]. Our analysis indicated that G6PD is ubiquitously expressed throughout the developmental stages and in adults (Figure 1B). IDH1 is highly expressed during early development, and the expression decreases and stabilizes 1 year after birth (Figure 1B). On the contrary, ME1 expression increases at 4 months post-conception week (PCW) and stabilizes thereafter (Figure 1B). Further, utilizing the human protein atlas (HPA) RNA sequencing database, we analyzed the expression of these genes in different brain regions, which also suggested that G6PD is highly expressed as compared to IDH1 and ME1 (Figure 1C) [31]. However, analysis of proteomics data from the human protein atlas (HPA) showed tissue-wise heterogeneity in the expression of these 3 genes (Figure 1D) [32]. Considering the differential expression of these genes and their basal level of expression across the brain tissues and other tissues indicates supplementation of citric acid and malic acid could increase the production of NAPDH in cells under G6PD deficient (G6PDd) condition. To validate our hypothesis, spectrophotometric NADPH estimation was performed by WST-8 assay, which indicated an increase in NADPH production in the human G6PD wild-type (G6PDwt) microglia cells treated with citric acid and malic acid (Figure 2E). We also determine the efficacy of citric and malic acid in reducing oxidative stress by flow cytometry. Our data indicated that citric and malic acid reduce the oxidative stress in G6PDwt microglia pre-activated with Lipopolysaccharides (LPS) (Figure 1F). There, both metabolic supplementation strategies can be used as potential therapeutics.

**Figure 1:**
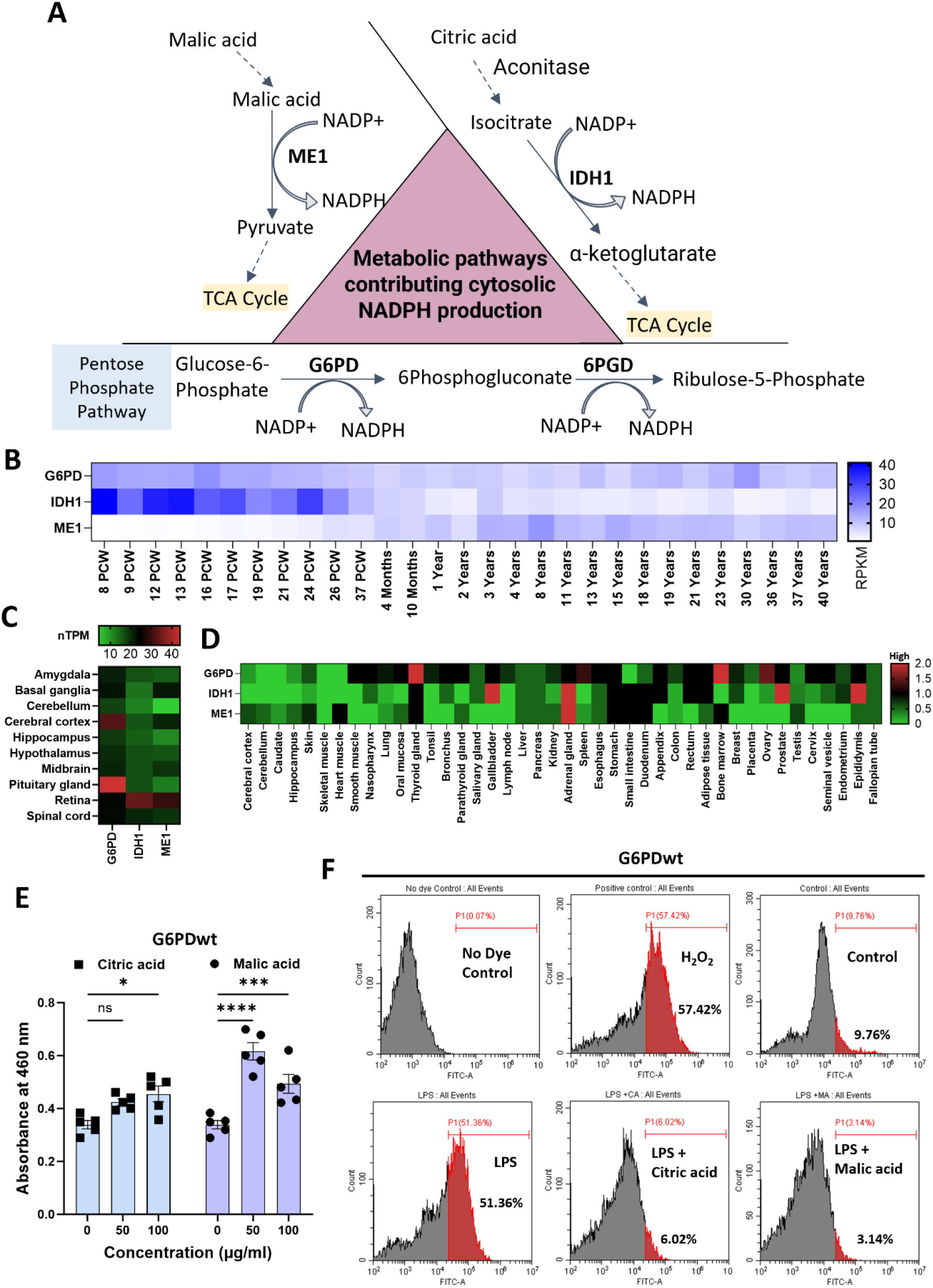
Expression analysis of cytosolic NADPH-producing genes for development and validation of therapeutic strategies: **A)** Diagrammatic representation of the pathways contributing to cytosolic NADPH pool; **B)** Heatmap represents FPKM (Fragments Per Kilobase of transcript per Million) value of the expression of cytoplasmic NADPH-producing genes from prenatal to 40 years adult brain RNA sequencing data from Brainspan database; **C)** Heatmap represents n-TPM (normalized Transcripts per million) values of expression of NADPH-producing genes in different brain regions derived from Human protein atlas database; **D)** Heatmap represents protein expression (low to high) of NADPH producing enzymes across all tissue types in human-generated by human protein atlas database; **E)** Bar graph represents absorbance (at 460nm) of WST8-NADPH estimation assay in microglia cells incubated with various concentrations of citric acid and malic acid. Data represent the mean ± SEM (n= 5). *Represents significance values compared to control, (**P=0.0022, ****p<0.0001). Statistical significance was calculated using ordinary two-way ANOVA; **F)** Histogram represents the percentage of ROS-positive microglia treated with LPS and LPS + metabolites. A gate on the histogram was placed while considering no dye control sample;

**Figure 2:**
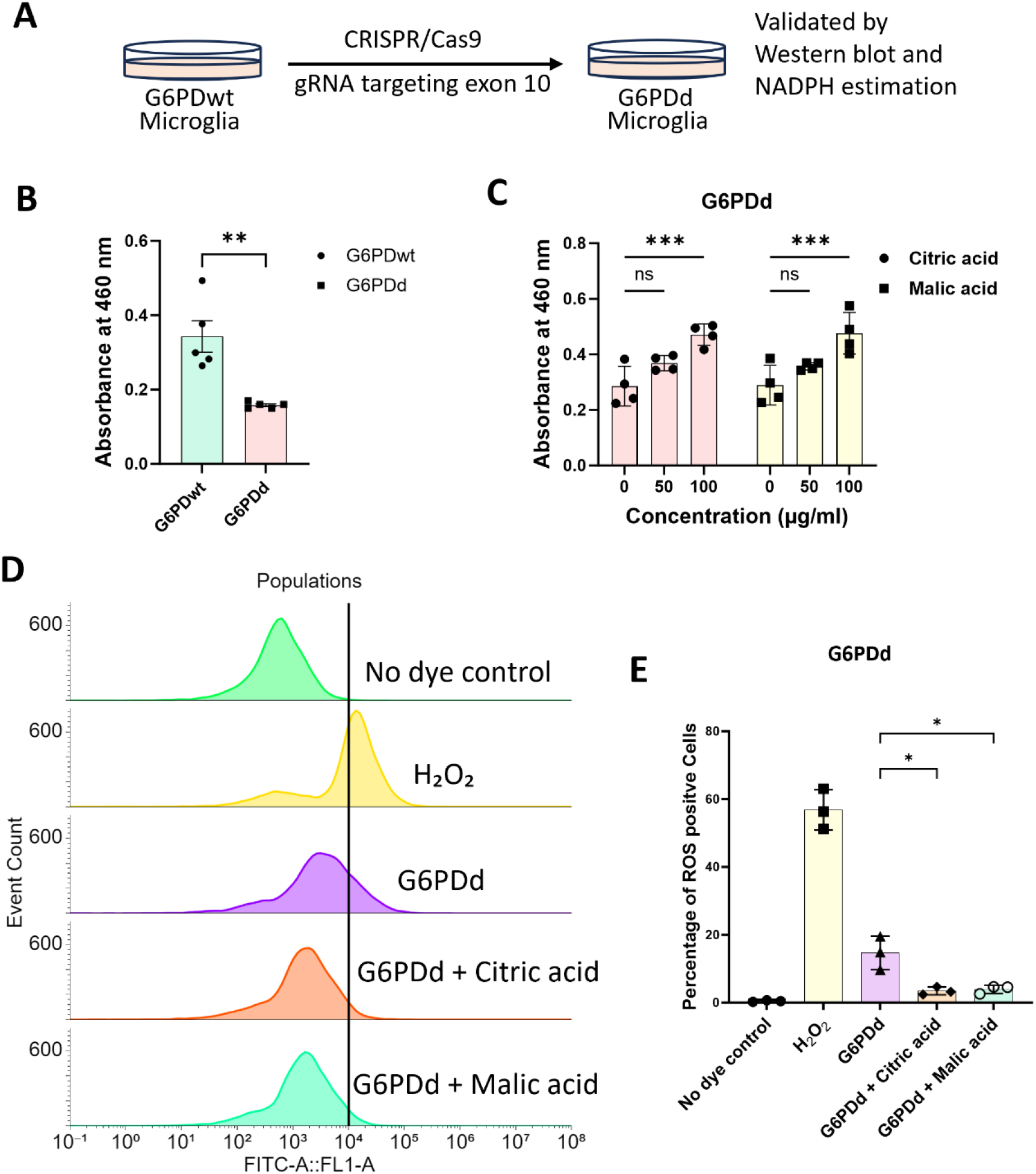
Metabolic supplementation increases NADPH production and reduces oxidative stress in G6PD-deficient microglia: **A)** Schematic representation of the generation of G6PD-deficient microglia. **B)** Bar plot represents WST-8 -NADPH estimation in G6PDwt versus G6PDd cells. Data represents the mean ± SEM (n=4). Statistical significance was calculated by unpaired t-test (**p=0.0024); **C)** Bar diagram represents absorbance at 460nm of NADPH estimation in G6PDd microglia at various citric acid and malic acid concentrations. Significance was calculated by two-way ANOVA (p= 0.0001); **D)** and **E)** Histogram and bar plot represent flow cytometry data of the percentage of ROS positive G6PD deficient microglia treated with LPS and LPS + metabolites; Statistical significance was calculated by two-way ANOVA (**p=0.0013).

### 3.2 Citric acid and malic acid increase NADPH production and reduce oxidative stress in G6PD-deficient microglia cells

To study the efficacy of the citrate and malic acid supplementation, CRISPR-mediated G6PDd microglia cells were used, which were previously generated by guide RNA targeting exon 10 of G6PD (Figure 2A) [28]. While characterizing the G6PDd microglia, we found a 50 - 60% reduction in NADPH level compared to G6PDwt. Thus, NADPH estimation was again performed in G6PDwt and G6PDd cells to confirm a reduction in level NADPH in G6PDd microglia for further use in therapeutics strategies (Figure 2B). Notably, G6PDd microglia supplemented with citric and malic acid exhibited an increase in NADPH production (Figure 2C). However, our previous study showed that G6PD deficient microglia exhibit increased basal oxidative stress as compared to wild-type [28]. Therefore, to check the efficacy of our metabolites in regulating oxidative stress, we performed flow cytometry analysis to capture the level of ROS in G6PDd microglia supplemented with metabolites. Our data indicated that supplementation of citric acid and malic acid reduces the percentage of ROS-positive G6PDd microglia from 20% to 4.5% (Figure 2D, E), which suggests their efficacy in regulating oxidative stress *in-vitro*.

### 3.3 Dieckol and resveratrol increase the expression of IDH1 and ME1 by nuclear translocation of NRF2

Recent reports also suggested that plant-derived small molecules (phytochemical) such as dieckol and resveratrol inhibit KEAP1 and allow nuclear translocation of NRF2 [33–38]. Therefore, we used dieckol and resveratrol to target the KEAP1-NRF2 pathway for increased expression of NRF2 downstream genes such as IDH1 and ME1 (Figure 3A). Initially, we determined the percentage of cell viability by MTT assay (Figure 3B). Dieckol within 20 μg/ml and resveratrol within 10μM concentration did not affect cell viability. Therefore, 10μg/ml (∼13.47μM) of dieckol and 0.5μM of resveratrol were used to develop therapeutics. We observed that the expression of IDH1 and ME1 was increased 1.5 to 2-fold (Figure 3C, D) after 24 hours of treatment with dieckol and resveratrol in G6PDwt and G6PDd microglia compared to untreated control. However, resveratrol-treated G6PDd microglia showed the highest expression of IDH1 and ME1 compared to G6PDwt (Figure 3D). Further, our immunofluorescent study confirmed that these phytochemicals increased the nuclear localization of NRF2, thereby enhancing the downstream expression of IDH1 and ME1 (Figure 3E, F). Thus, our experiment suggested that dieckol and resveratrol can be used to increase the expression of NADPH-producing genes to overcome the differential expression of this gene across all cell types for the development of combination therapeutics.

**Figure 3:**
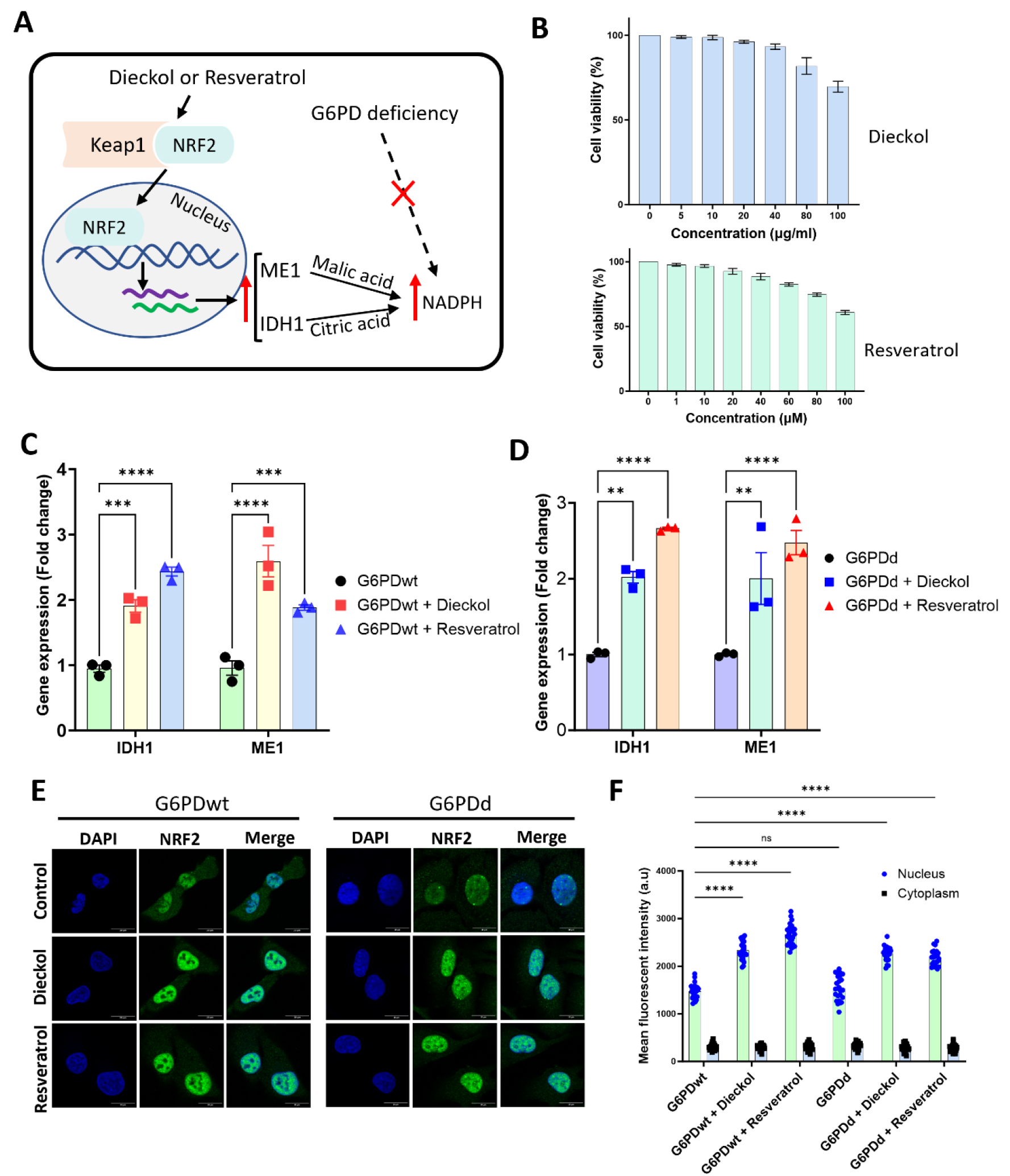
Phytochemical increases expression of IDH1 and ME1 in G6PDd microglia: **A)** Diagrammatic representation of dieckol and resveratrol-mediated nuclear translocation of NRF2 increases the expression of IDH1 and ME1; **B)** Line diagram showing MTT absorbance (at 570nm) value and respective IC_50_ of dieckol and resveratrol in microglia; **C)** and **D)** Representing qPCR fold change (2^-ΔΔCt) of gene expression of IDH1 and ME1 24 hours after treatment of dieckol and resveratrol in G6PDwt and G6PDd microglia (n=3). Two-way ANOVA was used to calculate the significance (***P=0.0002, ****p<0.0001 and **P=0.0013); **E)** Immunofluorescence image showing nuclear localization NRF2 in G6PDwt and G6PDd microglia treated with dieckol and resveratrol; **F)** Bar plot representing the mean intensity of nuclear versus cytoplasmic NRF2 level (n=15) in microglia treated with dieckol and resveratrol. Two-way ANOVA was performed to calculate the significance (****P<0.0001).

### 3.4 Combination therapeutics strategies increase NADPH production and restore redox balance and acidified lysosomes in G6PD deficient microglia

In our study, we also observed that dieckol and resveratrol upregulated the expression of IDH1 malic acid as combination therapeutics to increase treatment efficacy. Our data indicated that treating only dieckol and resveratrol to G6PDd microglia does not increase NADPH production (Figure 4A). However, the combination of small molecules along with metabolite increases the production of NADPH in G6PDd microglia (Figure 4A). Further, to evaluate the effectiveness of combination therapeutics on oxidative stress, we performed flow cytometry to detect the percentage of ROS-positive G6PDd microglia. Treatment of only citric acid or malic acid is sufficient to reduce the oxidative stress in G6PD deficient microglia *in-vitro* (Figure 4B). Additionally, the combination of dieckol or resveratrol with citric acid or malic acid showed a maximum reduction in the number of ROS-positive cells (1 - 3%) (Figure 4B). Our experimental data suggests that our phytochemicals and their combination with metabolites can be potential therapeutics for replenishing the NADPH pool and minimizing oxidative stress in the G6PDd microglia (Figure 4C). However, previously, we reported that the G6PD deficiency also affects lysosomal acidification in microglia [28]. Therefore, we also studied whether the supplementation of our therapeutics increased the number of acidified lysosomes. Our confocal microscopy data indicated an increase in lysosomal acidification (bright puncta of lysosomal vesicles) in all combinations of therapeutics (Figure 4C, D) compared to untreated control. Therefore, our *in-vitro* study revealed that the metabolic supplementation and its combination with plant-based small molecule NRF2 activator can effectively increase NADPH production, reduce oxidative stress, and restore lysosomal acidification, indicating their efficacy in treating G6PD deficiency.

**Figure 4:**
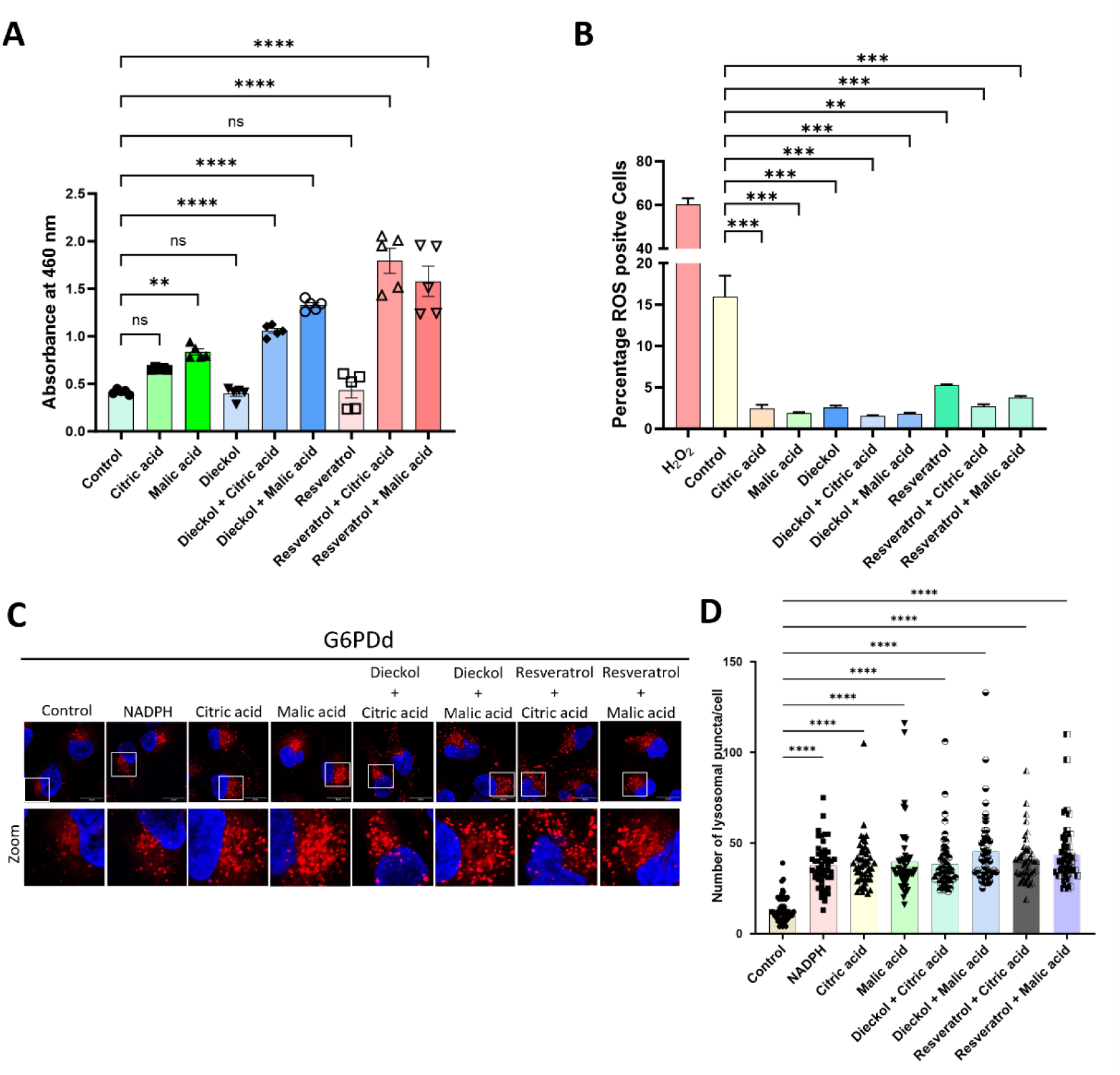
Combination therapy reduces oxidative stress and increases the number of acidified lysosomes in G6PDd microglia: **A)** Bar plot represents absorbance data of WST-8 -NADPH estimation assay in G6PDd microglia treated with various experimental conditions. Data (n=5) represented in mean ± SEM. Significance was calculated using one-way ANOVA (**P= 0.0069, ****p<0.0001, ns= 0.23 and ns >0.999); **B)** Bar plot representing a population of G6PDd microglia which are positive for ROS while treated with our therapeutics as compared to untreated control. Data represented here as mean ± SEM (n=3). Statistical significance was calculated by one-way ANOVA (**p=0.0019, ***p=0.0003, ****p<0.0001); **C)** The confocal microscopy images showing the number of acidified lysosomal vesicles in G6PDd microglia with and without treatment of our therapeutics. Corresponding zoom image exhibiting bright lysosomal puncta; **D)** Bar graph representing the number of acidified lysosomal puncta per G6PDd microglia (n=50) in our different experimental conditions. Significance was calculated using one-way ANOVA (****p<0.0001).

## 4. Discussion

A large cohort study has yet to elucidate the spectrum of G6PD deficiency disorders. Recently, G6PD deficiency has been linked with cardiovascular, diabetes, neurological, and autoimmune disorders [39–44]. Previously, we found that G6PD deficient microglia cells are insensitive to LPS-mediated ROS generation, which may play beneficial and detrimental roles during foreign pathogen invasion (ref). The high susceptibility of mutation in this gene is the major concerning factor for future disease burden globally. Besides, G6PD-deficient patients are sensitive to multiple drugs [7]. Therefore, patients should be screened for G6PD deficiency before prescribing drugs for certain diseases. Notably, no such medication is available to treat the G6PD deficiency. Earlier, clinical case studies using N-acetyl cysteine, α-tocopherol, and α-lipoic acid as therapeutics showed no promising results against G6PD deficiency [45]. In this study, we found that metabolic supplementation increased the production of NADPH and could reduce oxidative stress in the G6PD-deficient microglia. However, negative feedback mechanisms of metabolic pathways and differential expression of metabolic enzymes could be the drawbacks of metabolic supplementation therapy. Additionally, tissue heterogeneity is a major problem in metabolic supplementation therapeutics. To improve the efficacy of metabolic supplementation strategies, we planned to increase the expression of these genes using natural compounds targeting transcription factors regulating the IDH1 and ME1 expression. The previous study indicated that the KEAP1-NRF2 pathway is the primary regulator of cytoprotective responses to oxidative stress. KEAP1 is an adapter for cullin3 E3 ligase, which binds to the nuclear factor erythroid 2-related factor 2 (NRF2) and helps its degradation [46]. Generally, oxidative stress changes the conformation Keap1 and induces the release of NRF2 [47], which can further enter into the nucleus and upregulate genes essential for redox regulation, such as G6PD, heme oxygenase 1 (HO1) and glutathione peroxidases (GPXs) [48]. Previous studies also suggested that NRF-2 acts as a transcription factor for redox regulatory genes as well as IDH1 and ME1 [49,50]. Small molecule natural compounds nutraceuticals such as dieckol and resveratrol were previously shown to activate NRF2-mediated upregulation of redox regulatory genes, thereby decreasing oxidative stress in various disorders [51–54]. To circumvent tissue heterogeneity, we used a combination of therapeutic approaches to target the KEAP1-NRF2 pathway to increase the expression of IDH1 and ME1, followed by supplementation of citric and malic acid for enhancing the production of NADPH in G6PD deficient condition. Our *in-vitro* study of combination nutraceuticals using human G6PD deficient microglia showed promising results on NADPH production, regulation of oxidative stress, and lysosomal acidification. However, animal experiments should be conducted to conclude further the efficacy of combination therapeutics in G6PD deficient conditions.

## Acknowledgment

Ph.D. students AM and SM are grateful for their research fellowships to the Shiv Nadar Institute of Eminence (SNIoE), Delhi NCR, and the Shiv Nadar Foundation. Ph.D. student PU is grateful to the University Grant Commission (UGC) for the fellowship. Dr Soumya Pati is thankful to the Department of Science and Technology (DST/CSRI/2018/247). Prof. Shailja Singh is acknowledging Drug and Pharmaceuticals Research Programme (DPRP) (Project No. P/569/2016-1/TDT) and DST-SERB (Project No. CRG/2019/002231). We acknowledge the DST-FIST grant [SR/FST/LS-1/2017/59(c)] for the confocal microscopy facility at Shiv Nadar Institute of Eminence, Delhi NCR. We also acknowledge the Shiv Nadar Foundation Core Research grant.

## Authors contribution

Conceptualization: AM, SS and SP; Investigation and methodology: AM, IS, SM, and PU; Data analysis and Manuscript writing: AM; Manuscript Editing: SS and SP; Funding acquisition and supervision: SS and SP; All authors have read and agreed to the final version of the manuscript.

## Data availability

Data will be made available on request.

